## Supplementary figure for "Ecological drivers of CRISPR immune systems"

May 16, 2024

Wei Xiao,, University of Maryland

JL Weissman,, The City College of New York

Philip L.F. Johnson,, University of Maryland

### Supplementary Figures

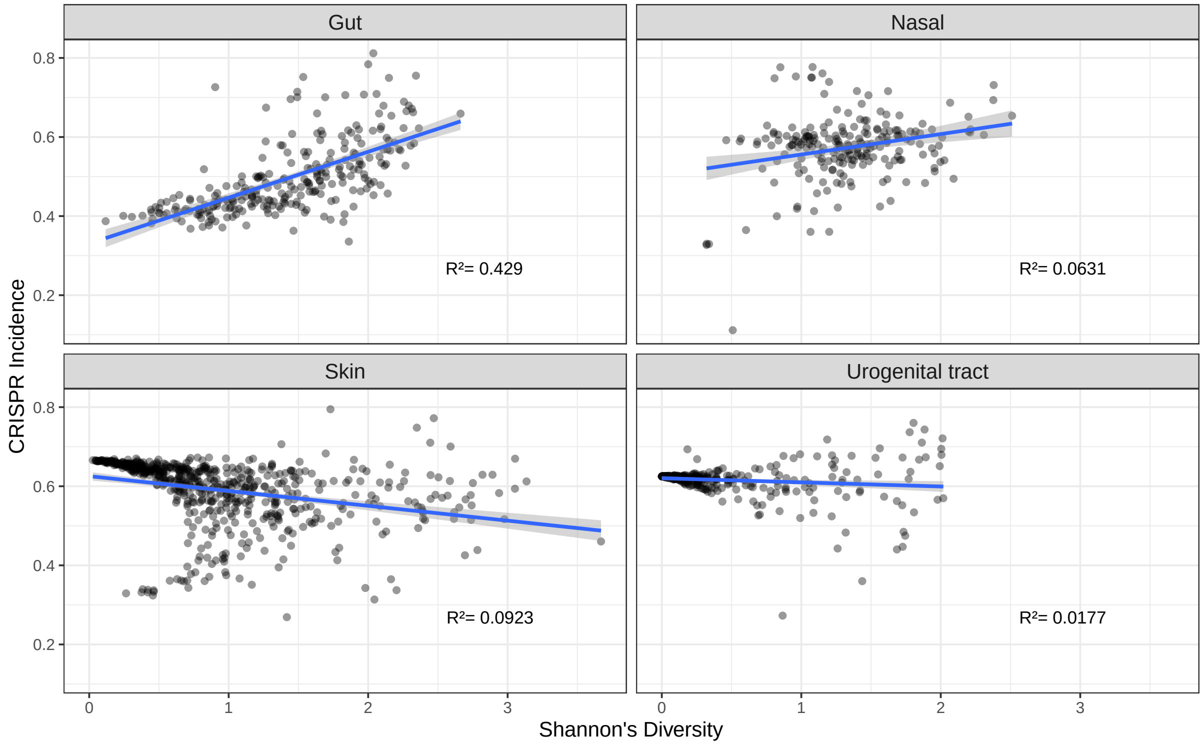


Figure S1: Correlations between CRISPR system incidence and microbial diversity vary across human body sites. The analysis was performed with HMP samples from gut, nasal, skin, and urogenital tract. Shannon's diversity index for these 1452 HMP samples was calculated with 16S rRNA sequencing data grouped at the genus level. CRISPR incidence came from CRISPRCasdb.

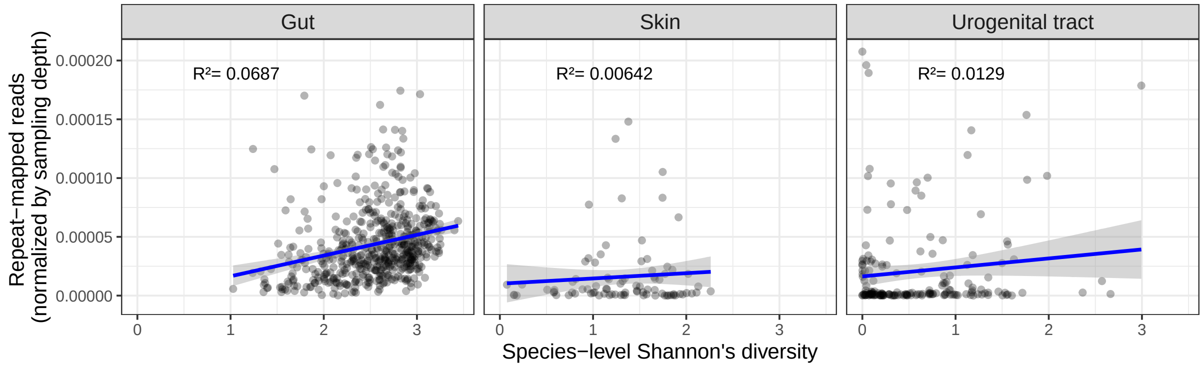


Figure S2: Correlations between sample-wise repeat-mapped read counts and Shannon's diversity index vary across human body sites. The analysis was performed with HMP metagenomic samples from gut, skin, and urogenital tract.

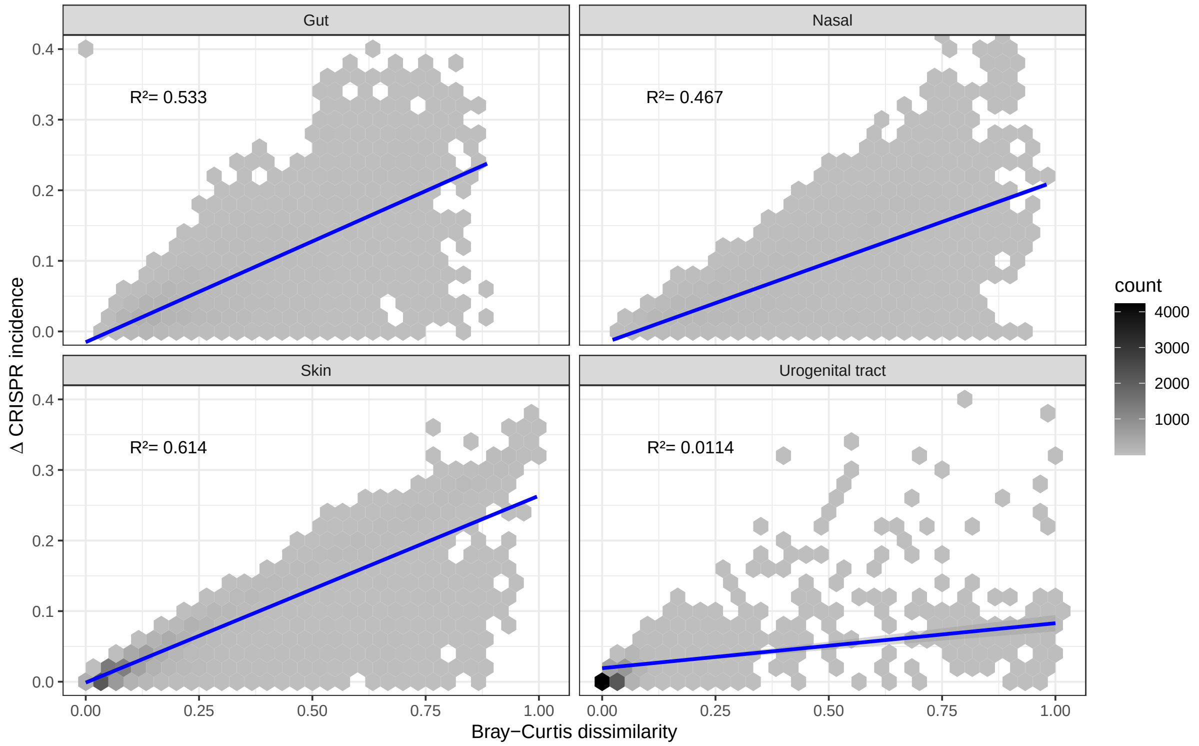


Figure S3: Correlations between the difference in CRISPR incidence between pairs of HMP 16S samples with similar $\alpha$-diversity (Shannon's index within 5% of each other) and the Bray-Curtis dissimilarity vary across human body sites. The analysis was performed with HMP metagenomic samples from oral, gut, skin and urogenital tract.

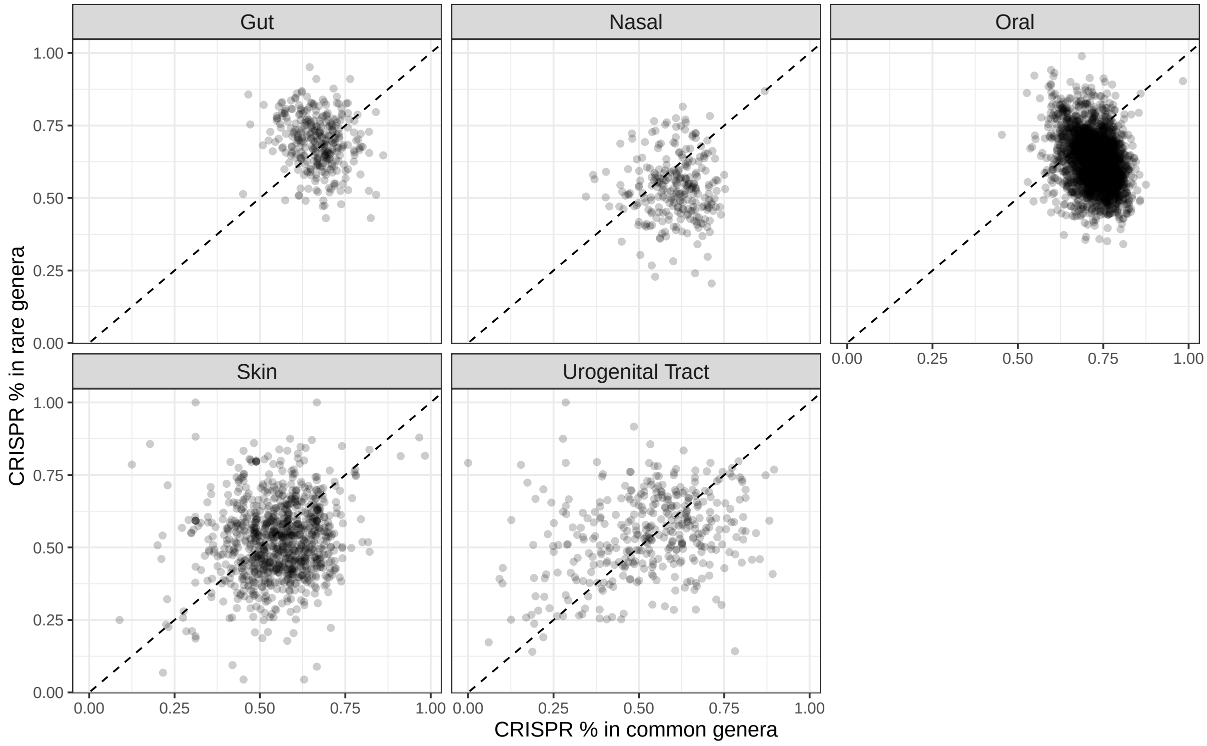


Figure S4: Within a HMP sample, CRISPR is found at a similar rate in rare genera (bottom 50% of the sample) and common genera (top 50% of the sample). Each point represents a single sample from HMP. Density analysis was performed with 16S rRNA data.

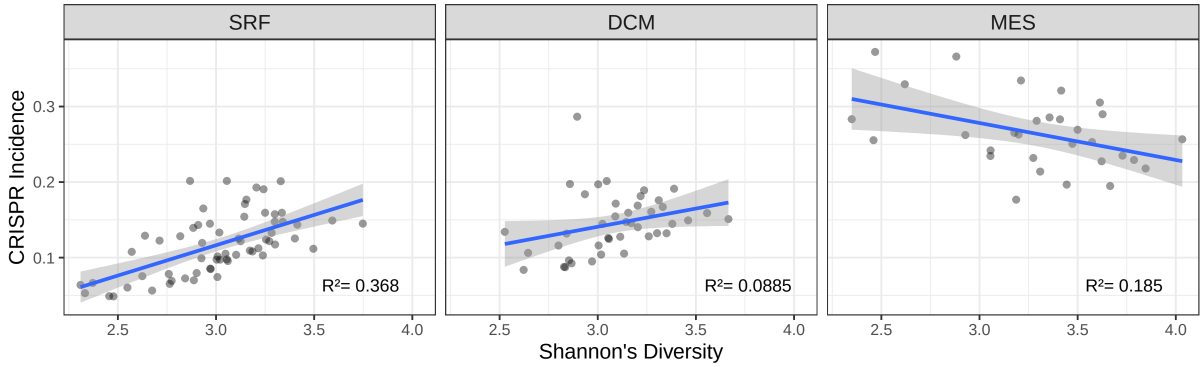


Figure S5: CRISPR system incidence shows various relationships with microbial diversity in ocean environments (SRF: $r^2 = 0.37$, DCM: $r^2 = 0.088$, MES: $r^2 = 0.18$). The analysis was performed with 135 the *Tara* Oceans samples of all 3 depths. The Shannon's diversity index of each sample was calculated with 16S rRNA sequencing data that were grouped at the genus level. The CRISPR incidence was annotated from all prokaryotes complete genomes and chromosomes of GenBank.

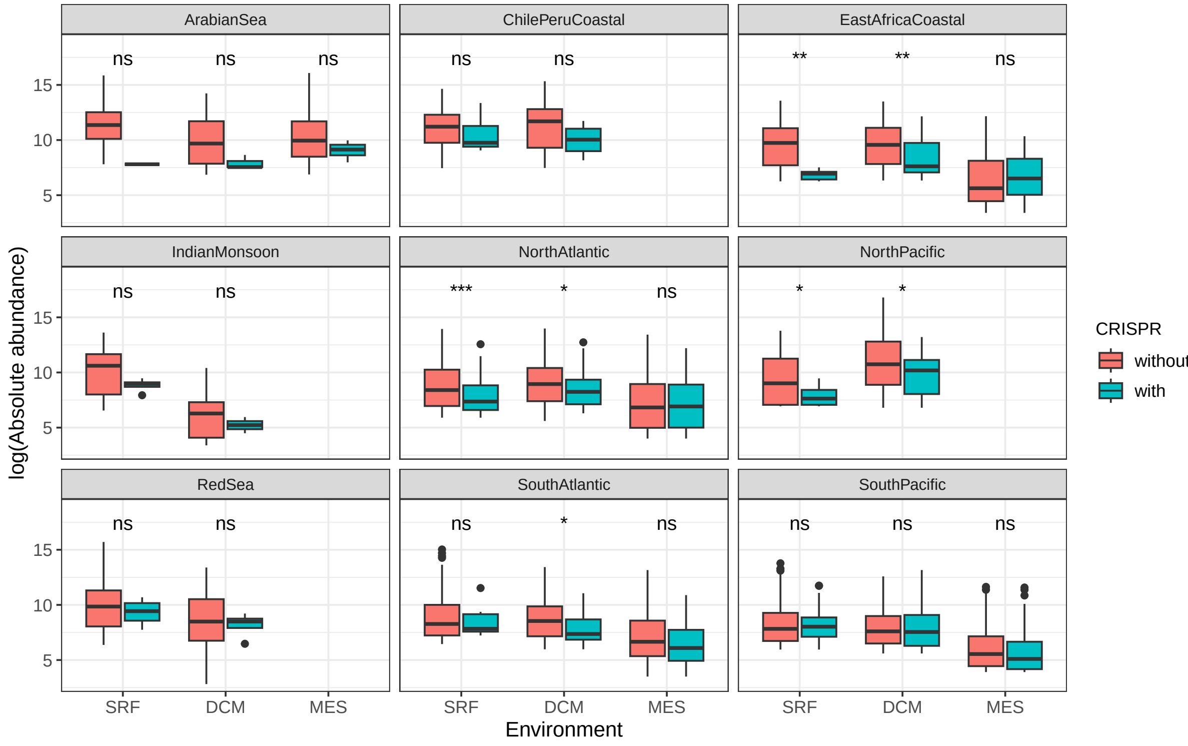


Figure S6: The figure shows the abundance of MAGs with and without CRISPR array(s) in different ocean sections of the *Tara* Oceans project. At DCM and SRF depths of the most of these areas, MAGs without CRISPR array(s) are more abundant than MAGs with CRISPR array(s).

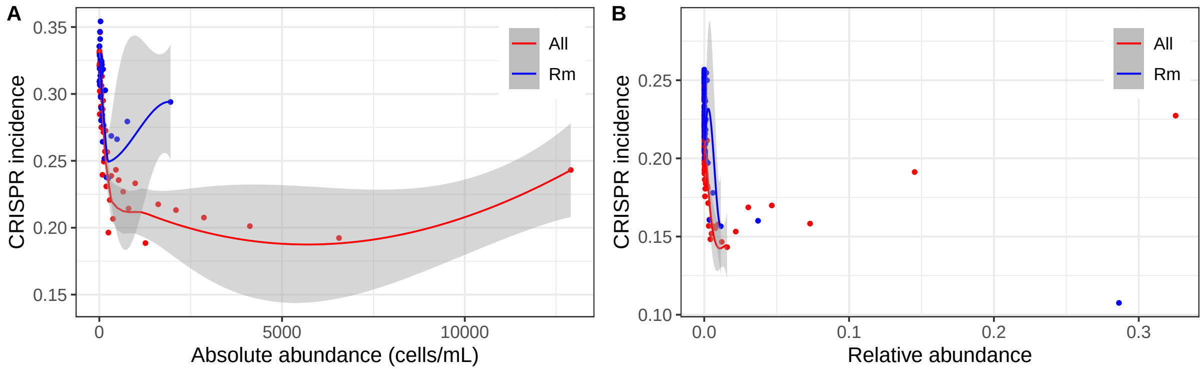


Figure S7: The negative correlation between CRISPR incidence and absolute abundance was driven by both prevalent and rare genera. Each point represents the median CRISPR incidence of 500 genera with a similar abundance of the *Tara* Oceans Project (A) and EMP (B). Red points are summarized data with all genera, while blue points are summarized data without prevalent genera (genera present in more than 75% of samples).

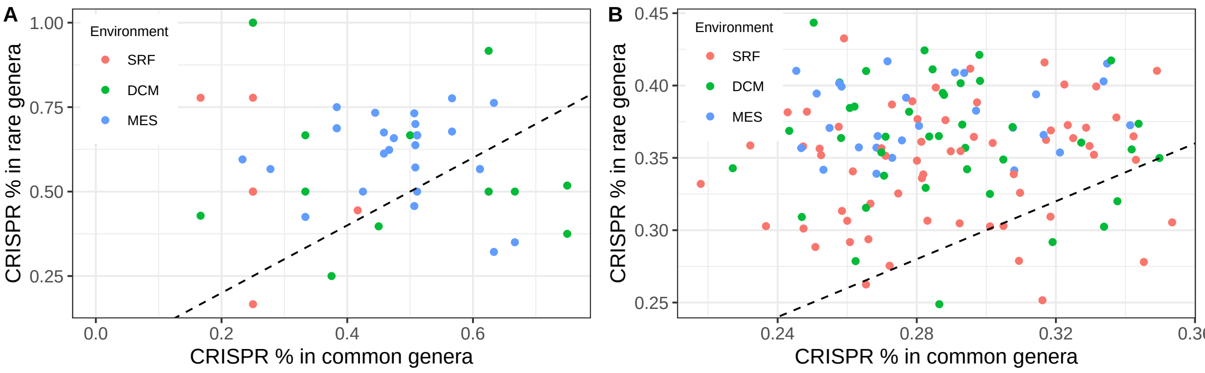


Figure S8: Lower density of archaea (A) or bacteria (B) associates with CRISPR incidence. Density was inferred from 16S rRNA data. Within a sample, CRISPR is found at a higher rate in rare archaea or bacteria genera (bottom 50% of abundances) than common archaea or bacteria genera (top 50% of abundances). Each point represents a single sample from the *Tara* Oceans Project. Points are colored by different sampling depth. SRF: surface water layer, ~-5m; DCM: deep chlorophyll maximum layer, ~-71m; MES: mesopelagic zone, ~-600m.
